# Post-perturbational transcriptional signatures of cancer cell line vulnerabilities

**DOI:** 10.1101/2020.03.04.976217

**Authors:** Andrew Jones, Aviad Tsherniak, James M. McFarland

**Author notes:** Co-senior authors (Aviad Tsherniak, James M. McFarland).

## Abstract

While chemical and genetic viability screens in cancer cell lines have identified many promising cancer vulnerabilities, simple univariate readouts of cell proliferation fail to capture the complex cellular responses to perturbations. Complementarily, gene expression profiling offers an information-rich measure of cell state that can provide a more detailed account of cellular responses to perturbations. Relatively little is known, however, about the relationship between transcriptional responses to perturbations and the long-term cell viability effects of those perturbations. To address this question, we integrated thousands of post-perturbational transcriptional profiles from the Connectivity Map with large-scale screens of cancer cell lines’ viability response to genetic and chemical perturbations. This analysis revealed a generalized transcriptional signature associated with reduced viability across perturbations, which was consistent across post-perturbation time-points, perturbation types, and viability datasets. At a more granular level, we lay out the landscape of treatment-specific expression-viability relationships across a broad panel of drugs and genetic reagents, and we demonstrate that these post-perturbational expression signatures can be used to infer long-term viability. Together, these results help unmask the transcriptional changes that are associated with perturbation-induced viability loss in cancer cell lines.

## Introduction

Large-scale efforts to map the chemical and genetic vulnerabilities of cancer cell lines have provided insight into cancer cell biology, gene function, and potential therapeutic avenues [25,8,28,2,27]. While these high-throughput experiments have relied on measuring changes in cell growth, there is a need to better understand the underlying shifts in cellular mechanisms that ultimately lead to a change in the growth phenotype.

In parallel efforts, large-scale databases profiling transcriptional responses to chemical and genetic reagents have opened the door for a deeper understanding of these effects by providing an information-rich measure of cellular response to perturbation [23]. However, the relationship between short-term post-perturbation transcriptional changes and longer-term changes in cell viability is not well understood.

A more detailed portrait of the transcriptional changes associated with viability loss would not only offer insight into fundamental modes of cancer cell death, but could also guide the development of therapeutic strategies. Several efforts have attempted to leverage transcriptional profiles to discover drugs’ mechanisms of action [14,21,23], but directly linking transcription and viability could expedite the discovery of promising drug-response mechanisms that could suggest therapeutic strategies that elicit one or multiple of these mechanisms. Furthermore, if early readouts of post-perturbation transcription are predictive of long-term changes in cell death, these transcriptional profiles could provide pharmacodynamic biomarkers that are indicative of a sample’s eventual response to treatment.

A number of studies have presented broad characterizations of the expression-viability relationship by examining responses to drugs with various mechanisms, revealing patterns in gene expression that are associated with fitness loss [18,24]. Another line of work has focused on building predictive models to reliably estimate a cell’s future viability response given the transcriptional profile [20]. While these studies have provided beneficial insights for a handful of compounds and cell contexts, there remains a need for a detailed characterization of the landscape of viability-related transcriptional responses across many chemical and genetic reagents.

Here, we investigate the relationship between transcriptional changes and cell viability following perturbation in cancer cell lines by integrating several large-scale datasets, and applying simple and interpretable statistical analyses. We find transcriptional “signatures” associated with loss of viability that are robust and biologically meaningful for chemical and genetic perturbations, and enable stratification of samples based on inferred sensitivity to a given perturbation, which is a key goal of developing targeted therapeutics. Furthermore, we observe clusters of perturbations exhibiting similar expression-viability relationships, which could aid the identification and refinement of perturbation classes. These analyses not only enhance our understanding of pathway regulation induced by a variety of treatments, but also provide an avenue for using short-term transcriptional readouts for predicting long-term sensitivity to these treatments.

## Results

### A global transcriptional signature for sensitivity to small molecules

We first sought to identify the changes in gene expression that are associated with drug-induced loss of long-term cell viability across a large set of compounds and cell lines. To this end, we leveraged data generated by the Connectivity Map using the L1000 assay, which measures the expression levels of 978 landmark genes following perturbation and computationally infers the remainder of the transcriptome. In parallel, we integrated drug sensitivity data from three sources: the PRISM Repurposing dataset [3], the Genomics of Drug Sensitivity in Cancer resource (GDSC) [8,1,21,22,11], and the Cancer Target Discovery and Development database (CTD2) [22,1,21] (Methods). Together, these viability datasets have screened thousands of small molecules in hundreds of cell lines. We integrated the expression and drug response datasets by pairing profiles from each dataset with the same perturbation, cell line, and sufficiently similar dose, yielding a large set of matched profiles, each of which consisted of a cell line’s differential expression following treatment with a specific compound, along with the sample’s observed viability following treatment. After selecting for compounds with a sufficient number of cell lines, this integration yielded over 12,000 transcriptional profiles spanning 147 compounds, 54 cell lines, and 2 time-points (6 and 24 hours). Importantly, the cell viability measurements were taken at 3-5 days after initial drug treatment (Methods).

We assessed the linear association between each gene’s transcriptional response and the measured viability effects of the perturbation, yielding a transcriptional signature associated with decreased cell viability across over 5,000 profiles (141 compounds) (we refer to this association as the “global viability signature”, as it considers the average association across all drugs and cell lines) (Figure 1A, Methods). This analysis revealed a robust association between drug sensitivity and expression for many genes, indicating that there is a consistent transcriptional signature related to viability effects. Specifically, the global viability signature showed enrichment for genes involved in pathways relevant to cell viability, such as cell cycle regulation and apoptosis, suggesting that this approach is able to identify the biological pathways associated with fitness loss across a broad range of drug classes and different cell types (Figure 1B, 1C).

**Fig. 1:**
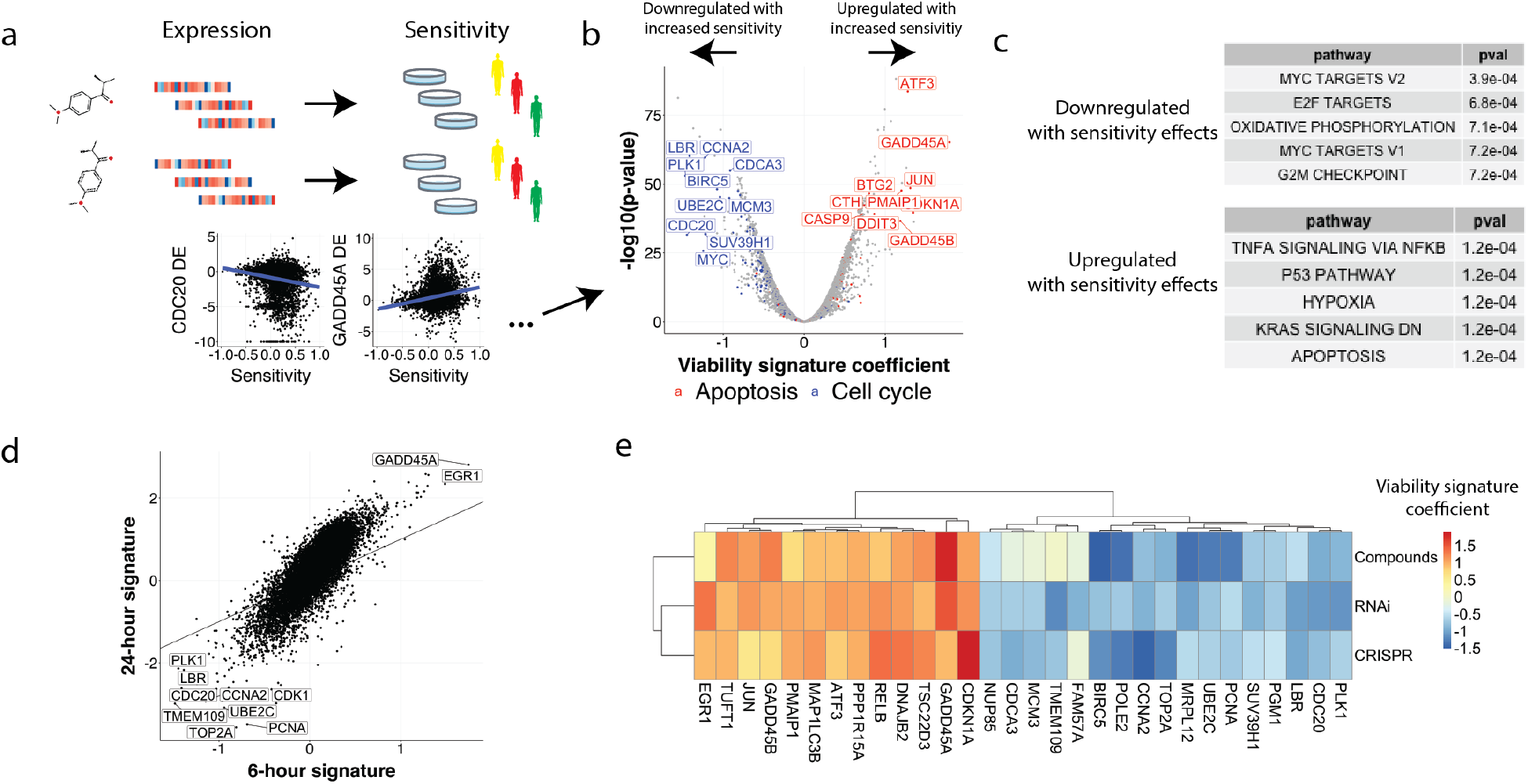
The global viability signature finds biologically interpretable associations between gene expression and viability. (a) Schematic showing integration of large-scale datasets of measurements of transcription and viability following treatment with a perturbations, which allows for understanding changes in expression that are associated with longer-term loss of viability. (b) We computed the global association between post-treatment change in expression and sensitivity, and the top genes in the global “viability signature” were involved in pathways related to cytotoxic processes, such as apoptosis and cell cycle regulation. (c) A gene set enrichment analysis of the global viability signature reinforced the enrichment of cytotoxic pathways, with cell cycle-related genes being downregulated and apoptosis-related genes showing upregulation in association with sensitivity. (d) Viability signatures for expression measurements at 6 and 24 hours post-treatment. A positive trend emerges, although the signature at 24 hours is stronger in magnitude. The solid line indicates unity. (e) Heatmap showing the association between change in gene expression and sensitivity, computed separately for compounds, RNAi, and CRISPR. The gene-wise relationship between expression and viability remains consistent across perturbation technology types, as shown here with the genes with strongest average association across types.

### Global viability signature generalizes across perturbation types, time-points, and independent datasets

Having observed a global relationship between changes in transcription and cell viability across the large L1000 chemical perturbation dataset, we sought to determine whether the association remained consistent across perturbation types, experimental conditions, and independent datasets.

To test whether a similar transcription-viability relationship is induced by genetic perturbations, we leveraged data from the Broad Institute’s Cancer Dependency Map, which has measured the viability re-sponse of hundreds of cell lines in genome-wide RNAi knockdown and CRISPR/Cas9 knockout experiments [25,17,5,6]. We integrated the Connectivity Map’s RNAi and CRISPR/Cas9 transcriptional profiles with the associated viability measurements — here, matching profiles by cell line and targeted gene, together totaling nearly 10,000 transcriptional profiles covering 935 target genes and 76 cell lines — in order to identify a viability-related transcriptional signature.

The global viability signatures elicited by RNAi and CRISPR generally agreed with that identified for small molecules (Pearson R: compounds/RNAi = 0.61, compounds/CRISPR = 0.59, RNAi/CRISPR = 0.50) (Figure 1E), suggesting that, on average, the regulation of transcriptional pathways related to decreased cell fitness is consistent across different perturbation types.

Furthermore, the concordance of transcription-viability relationships across datasets suggests that the global viability signature estimated from small molecule perturbations was not specific to the composition of compound libraries used for screening. In other words, the global viability signature seems to generalize beyond the small molecules present in the chemical screening datasets. This point was further reinforced by the concordance of viability signatures estimated from three individual chemical sensitivity datasets, each of which was generated by an independent research group (Figure S1).

As transcriptional signatures of chemical perturbations have been shown to evolve over the course of hours to days, a key question is how the estimated viability signatures change over time. The Connectivity Map contains profiles at 6 and 24 hours post-perturbation for many compounds, so we assessed whether there were differences in the expression-viability relationships identified at each of these time points. We found that the estimated global signatures for the two time points were highly correlated (Pearson R = 0.82), indicating that the expression-viability relationship remains largely consistent over this time period, and that patterns in expression associated with long-term viability effects could be detectable as early as 6 hours after perturbation (Figure 1D).

We also found that the association between transcriptional response and drug sensitivity was stronger in magnitude at 24 hours (Figure 1D). Interestingly, transcriptional responses for some genes showed strong association, as measured by the viability signature coefficients, with viability at 24 hours but virtually no association at 6 hours post-treatment, including CDC45 and CDK1, both of which are involved in cell cycle regulation and mediating response to DNA damage, indicating that the post-perturbation time course of different viability-related cellular processes may vary.

### Predicting viability from short-term transcriptional response

Having validated the robustness of the global expression-viability relationship, a key question is whether a cell line’s long-term drug sensitivity can be computationally predicted from short-term transcriptional changes. To test this, we trained and evaluated predictive models, using over 12,000 transcriptional profiles comprising data pooled across compounds and cell lines. We used the 978 landmark gene expression values as inputs to the models and restricted these analyses to compounds that were screened in at least 15 cell lines in order to ensure that we could robustly assess each model’s predictions on a per-compound basis (using 10-fold cross-validation; Methods).

The models showed moderate predictive accuracy overall (Pearson R between predicted and measured viability = 0.34), reinforcing the robustness of the association between expression and viability globally across compounds and cell lines (Figure 2A). For comparison, we also trained models using only a scalar-valued measure of the overall strength of expression changes (transcriptional activity score, TAS [23]), rather than the specific pattern of expression response. Surprisingly, TAS predicted sensitivity fairly well, suggesting that even simple readouts of the aggregate magnitude of transcriptional change could be predictive of sensitivity for many classes of compounds; however, models trained using the full, multivariate expression profiles showed modest, but consistent, improvement (Figure 2A).

**Fig. 2:**
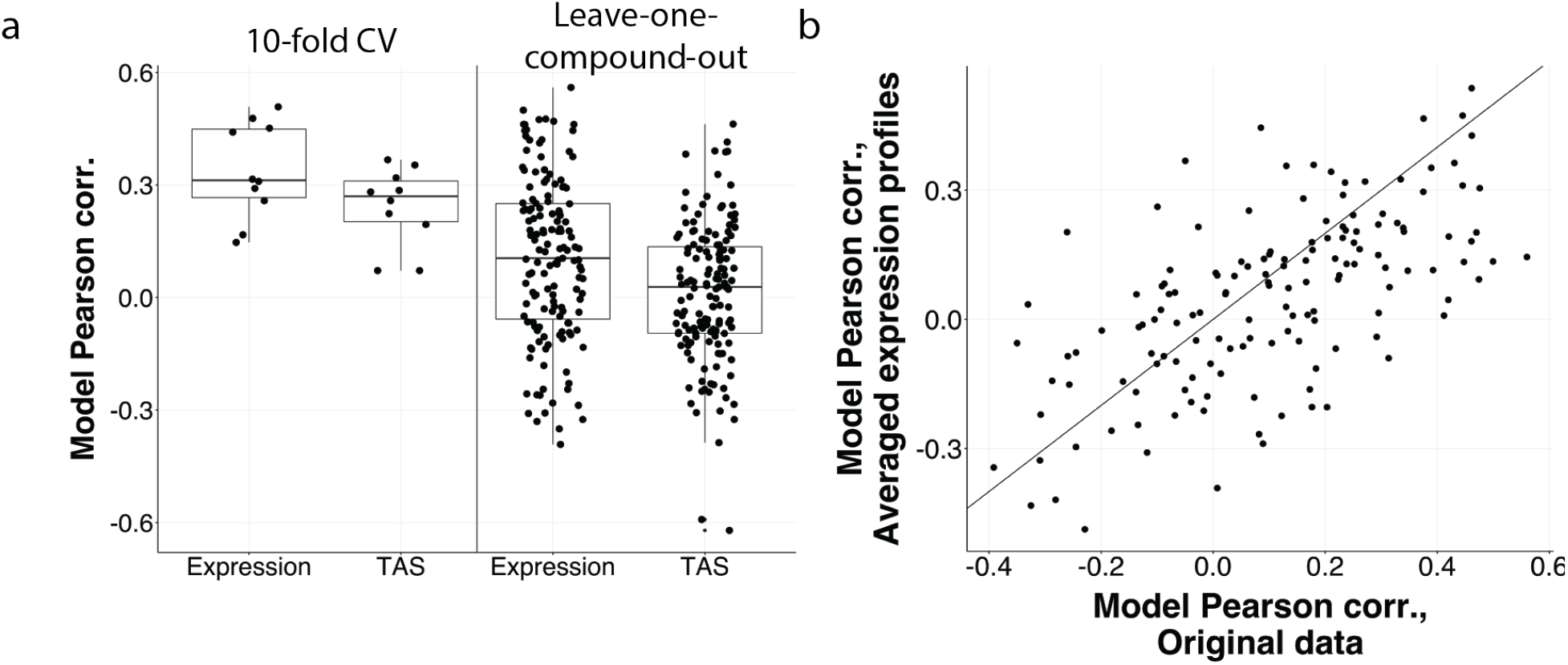
Predicting drug sensitivity using profiles of differential expression following chemical perturbation. (a) Left: Using transcriptional response profiles to predict sensitivity across compounds achieves moderate predictive performance using elastic net regression (left two boxes); however, predicting compound-specific sensitivity differences across cell lines yields a substantially larger range in performance, suggesting the model is primarily fitting and predicting differences among average compound responses. Each point on the left indicates the performance of one of 10 folds; each point on the right corresponds to the performance of a model for one compound. We also note that overall expression magnitude (TAS) also predicts viability fairly well. (b) Comparison of predictive models trained on all pooled transcriptional profiles (x-axis) and on profiles that have been mean-collapsed across cell lines for each compound. Each point represents the performance of the models for one compound. The comparable ability of the two models to predict sensitivity to individual compounds suggests that the global viability signature is primarily driven by variation between compounds.

While these models capture a consistent relationship between transcription and viability, they are measuring an average, or “global”, effect across compounds. Given the heterogeneity in the classes of small molecules in these datasets, it is important to investigate the nature of transcriptional effects that are specific to individual compounds or compound classes and how these relate to their viability effects on cancer cell lines.

To make a first approximation of the level of compound-specificity in the transcription-viability relationships, we tested whether a global viability signature (trained on data pooled across cell lines and compounds) could predict differential sensitivity across cell lines for individual held-out compounds (here, we only test on compounds which produce sufficient differential viability effects across cell lines; Methods). Notably, although the global transcriptional patterns predicted sensitivity better than TAS, these models showed considerable variability in performance across compounds when predicting selective killing of individual compounds (Figure 2A). This result suggests that, although the global expression-viability relationship can predict differential sensitivity between cell lines for some compounds, there exist substantial compound-specific differences in the expression-viability relationship.

Because the predictive models pooled all the data together for training, the expression signatures identified are necessarily shared among compounds, and will be driven by both differential sensitivity across cell lines to a given compound, as well as compound-to-compound differences in overall cell killing and expression responses. To tease apart these effects, we trained a model on expression profiles that had been averaged across cell lines for each compound (explicitly removing any cell-line-specific component of the transcriptional responses), and found that this model predicted sensitivity as well as the model trained on the original profiles (Figure 2B). This result suggests that the models are strongly influenced by variation across compounds, rather than differences across cell lines in the sensitivity to a given compound.

Although the global viability signature and these predictive model analyses demonstrate a transcription-viability relationship that is partially shared across compounds, it is vital to account for patterns that are unique to individual compounds or drug classes. Thus, we next sought to more precisely understand the extent to which each compound induces its own unique viability-related response.

### Assessing perturbation-specific viability signatures

Given the large variability across compounds in our ability to predict cell lines’ sensitivity with the global signature, we turned to directly assessing the transcription-viability relationship for each individual compound as a means to more precisely assess the level of compound-specificity in the expression-viability relationship.

In order to understand the relationship between transcriptional response and long-term viability effects for each compound, there is a need to disentangle changes that occur selectively in sensitive cell lines and those that occur in all cell lines. The former is important for identifying expression patterns of selective killing, while the latter is useful for understanding the baseline effects of a drug. For example, in the expression profiles following treatment with tanespimycin (an HSP inhibitor), some genes (e.g., PTK2) show a stronger change in expression in more sensitive cell lines, and other genes (e.g., HSPA6) show a change in expression even in insensitive cell lines (Figure 3B).

**Fig. 3:**
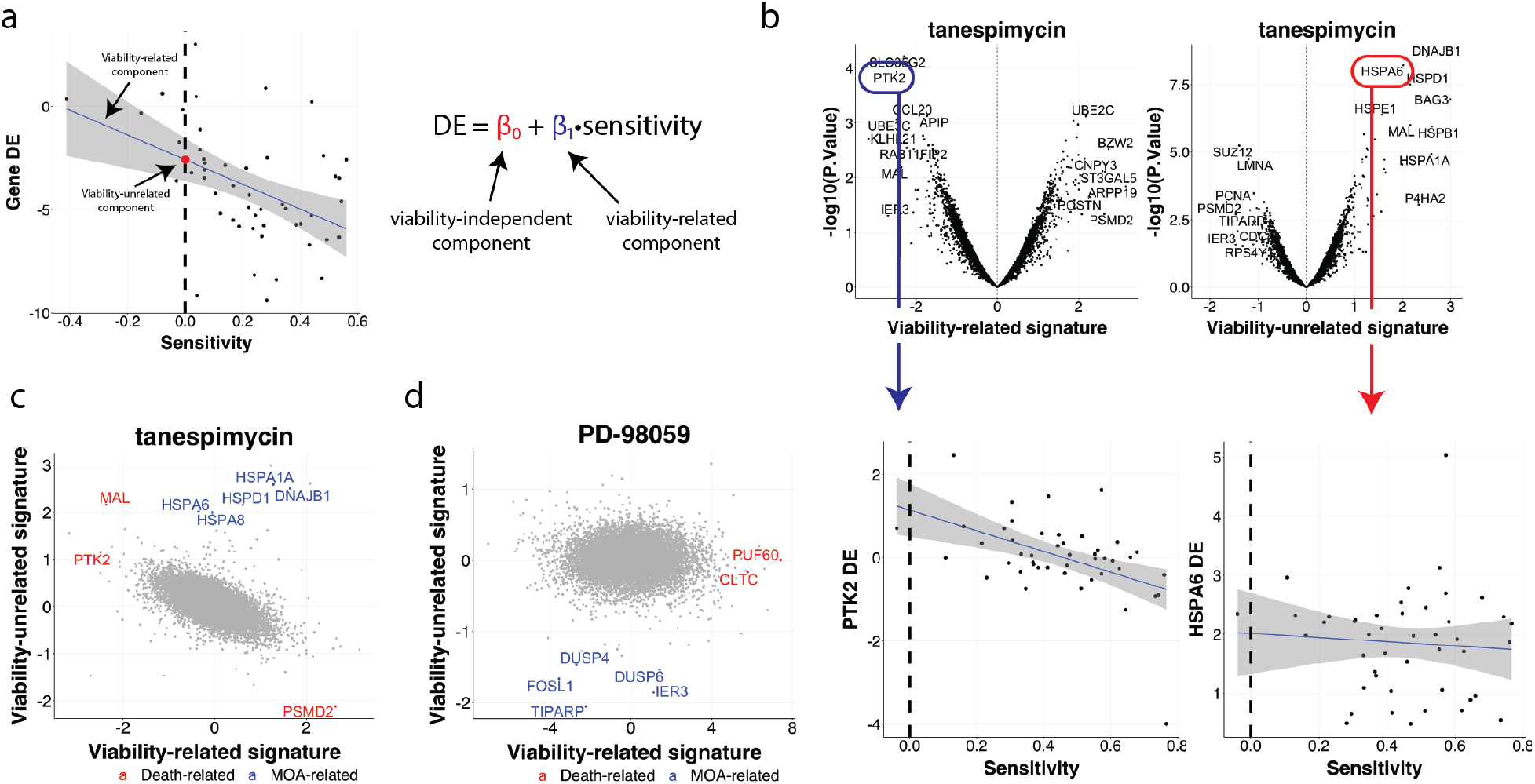
Compound-specific viability signatures reveal viability-related and -independent components of expression signatures. (a) An example relationship between sensitivity and differential gene expression for a single compound (each point is a cell line). We define viability-related and -independent components of the expression response as the slope and intercept of this line, respectively, which directly isolates the signature that is associated with loss of viability. (b) Viability-related (top left) and -independent (top right) signatures for tanespimycin, an HSP inhibitor. Tanespimycin induces a change in the expression of some genes (e.g., PTK2, bottom left) in association with the sensitivity to vemurafenib, while the expression of other genes (e.g., HSPA6, bottom right) changes regardless of the sensitivity. (c) Viability-related and -independent components recover death- and MOA-related signatures, respectively, for tanespimycin. (d) Same as (c), but for PD-98059, a MEK inhibitor.

To deconvolute these effects, we used linear models to account for these two components of the transcriptional response for each compound (Figure 3A): one that measures the expression-viability relationship (slope term), and another that measures changes in expression that occur independent of viability effects (intercept term). We refer to these as the “viability-related” and “viability-independent” signatures, respectively (Methods). This approach allows for a more precise dissection of the components of post-treatment expression that are directly related to eventual selective fitness effects.

For example, by estimating these signatures for tanespimycin, we found that the viability-related signature showed enrichment for genes related to cell death, such as MAL and PTK2 (Figure 3C), and a gene set enrichment analysis showed strong downregulation of gene sets related to cell cycle. On the other hand, the viability-independent component showed a strong representation of genes related to heat shock response and DNA damage, including HSPA1A, HSPD1, and DNAJB1, suggesting that the viability-independent component could be useful for isolating genes related to a compound’s mechanism of action.

By extending this approach across all compounds with a sufficient range of killing, we found that both the viability-related and -independent components exhibited biologically meaningful patterns for many drugs. For example, PD-98059 (a MEK inhibitor) showed a viability-related signature that was enriched for cytotoxicity-related genes (PUF60 and CLTC), while its viability-independent signature was enriched for MAPK pathway genes (DUSP4/6 and FOSL1) (Figure 3D).

Importantly, we found differences in viability-related signatures between compounds, reinforcing that a single signature cannot capture the diversity of compound-specific responses (Figure S2).

We also examined the analogous landscape of perturbation-specific viability signatures for genetic reagents using the same modeling framework. Because relatively few cell lines have been screened in the CRISPR L1000 data, we restricted our analysis to the RNAi data, composed of 268 unique target genes and 66 unique cell lines after filtering for perturbations profiled in at least 9 cell lines in order to be powered to estimate the variability across cell lines.

Similar to the compound-level signatures, we found that the gene-level viability signatures — computed on profiles averaged across shRNAs for each target gene — were enriched for genes related to cell death, and for pathways related to specific genes’ functions. For example, the viability-related signature associated with MAP2K1 knockdown in sensitive cell lines was enriched for genes related to apoptosis and MAPK signaling, such as IER3 and DUSP6, and shows concordance with the viability signature for the MEK-inhibitor selumetinib (Figure 4C).

**Fig. 4:**
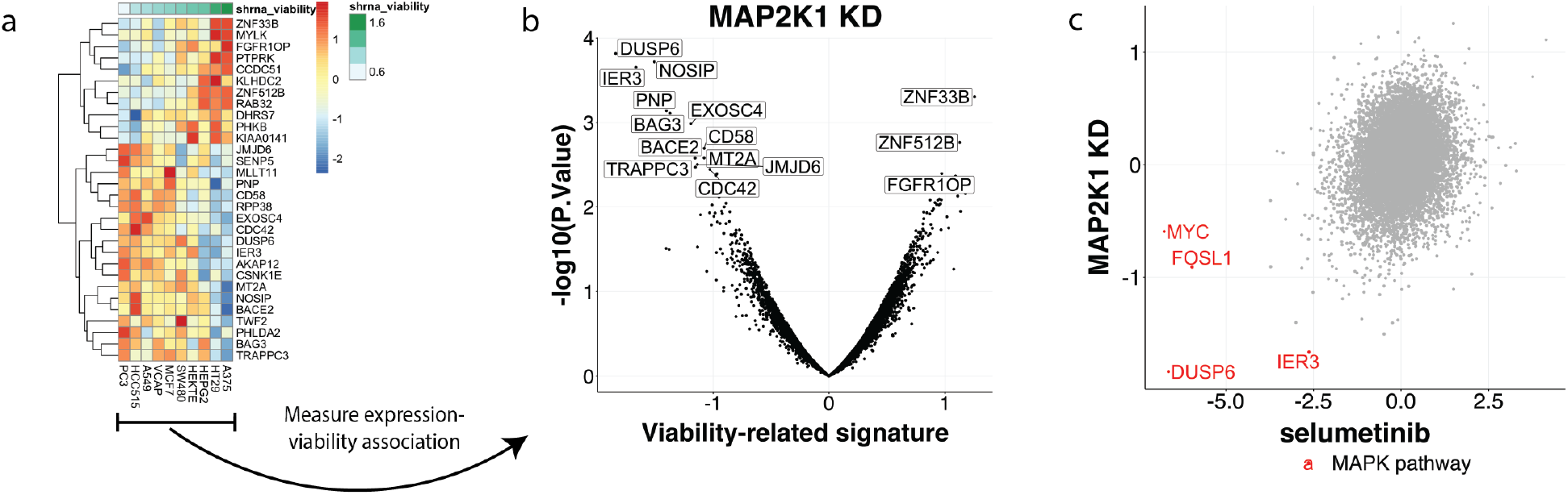
Gene-specific signatures for RNAi capture transcriptional response to genetic reagents associated with viability loss, and allow for making connections between genetic perturbations. (a) Differential expression following MAP2K1 knockdown showing strong expression response in cell lines which are sensitive to MAP2K1 inhibition. (b) Viability-related signature component of expression following MAP2K1 knockdown. (c) Comparison of viability-related signatures for MAP2K1 knockdown and selumetinib (a MEK inhibitor) shows a consistent implication of MAPK pathway-related genes in the expression-sensitivity association.

We again observed robust differences between viability-related signatures across targeted genes, demonstrating the heterogeneity of perturbation-specific expression-viability relationships. Together, these observations suggest that although there is consistency in viability-related transcriptional signatures among subsets of compounds, these signatures vary substantially across the full panel of perturbations analyzed here.

### Identifying clusters of viability-related signatures

Given the heterogeneity of expression-viability relationships across perturbations, we next tested whether certain groups of perturbations elicited similar patterns of viability-related signatures, and whether these coincide with more traditional mechanism-based classifications. To do so, we performed a joint hierarchical clustering of 415 perturbation-specific chemical and genetic viability-related signatures (Figure 5A).

**Fig. 5:**
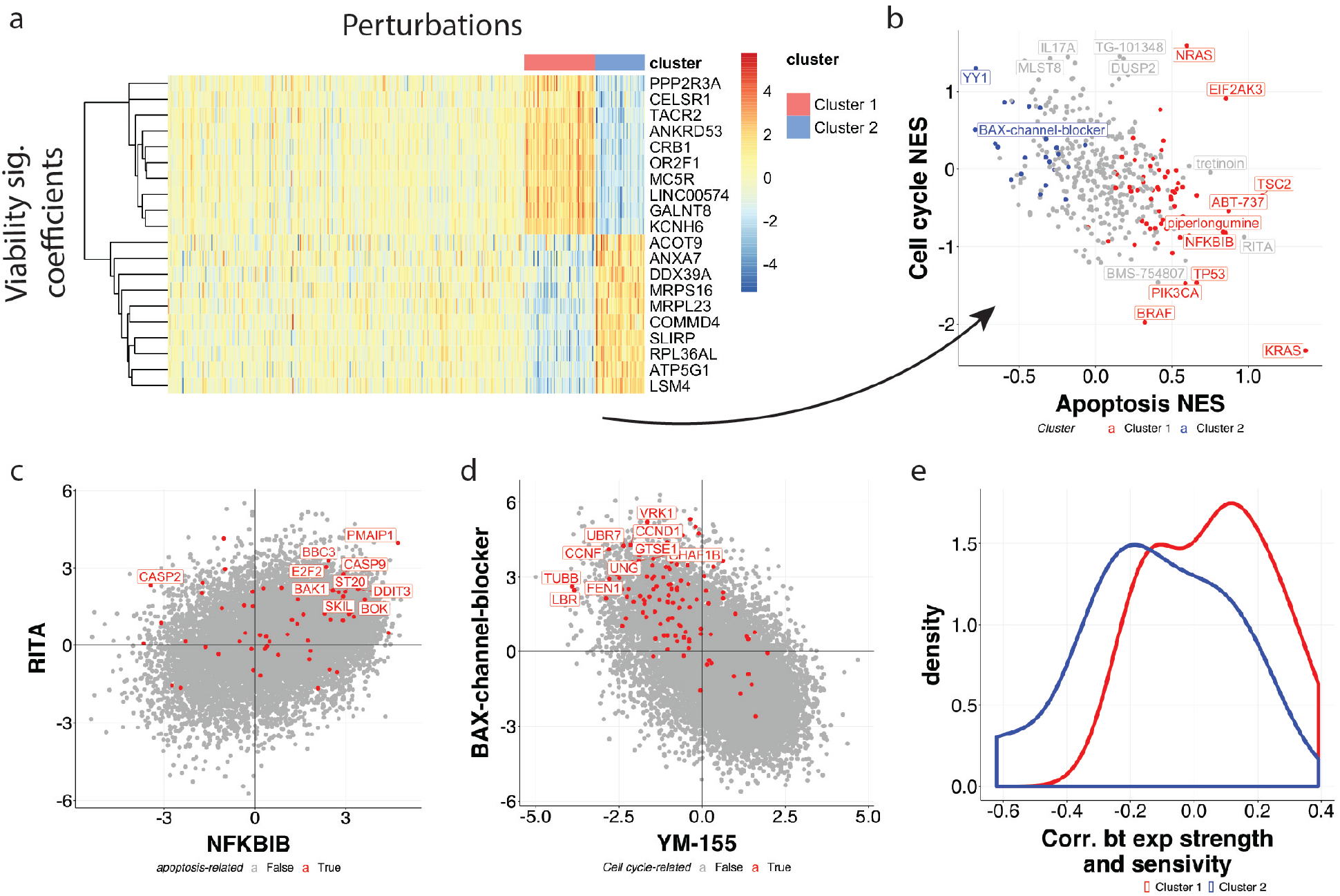
Joint clustering of chemical and genetic viability signatures reveals groups of perturbations with reversed expression-viability relationships. (a) Heatmap showing the perturbation-specific viability-related signatures for chemical and genetic perturbations for a set of top genes. Two clusters of perturbation emerge: cluster 1’s signatures resemble the global viability signature, and the perturbations in cluster 2 induce a distinct, somewhat anticorrelated transcriptional signature. Individual perturbation names are omitted for clarity (see Supplementary Material for full list). (b) Enrichment of cell cycle- and apoptosis-related genes for each perturbation’s viability-related signature, as measured by normalized enrichment scores (NES, Methods). The two clusters of compounds are particularly distinguishable based on their apoptosis-related effects. (c) The concordance between RITA (an MDM inhibitor) and NFKBIB (a gene implicated in apoptosis) viability-related signatures suggests that the viability signatures are able to detect interesting changes in pathways related to cell death. (d) Comparing the viability-related signatures for YM-155 (a survivin inhibitor) and BAX-channel-blocker (a cytochrome C release inhibitor) reveals a negative relationship, indicating that these compounds might have opposing effects on the expression-viability relationship. (e) Density plot showing the distribution of correlation coefficients between overall transcriptional response magnitude and sensitivity for the perturbations in both clusters. The density for cluster 1 suggests overall differential expression is greater in samples with higher sensitivity for this cluster’s perturbations, while the negative shift of cluster 2’s density implies that these compounds induce a stronger transcriptional change in samples with lower sensitivity or even heightened proliferation.

As expected, the clustering revealed substantial heterogeneity across perturbation-specific signatures. Several interesting connections among viability signatures within and between perturbation types emerged; for example, RITA (an MDM inhibitor) and NFKBIB knockdown yielded concordant viability-related signatures, indicating that the transcriptional responses associated with sensitivity to these perturbations were similar (Figure 5C).

More generally, this analysis revealed several clusters that spanned genetic and chemical perturbations. The predominant cluster — composed of compounds like teniposide and nutlin-3, and genetic perturbations like BCL2L1 and MCL1 — consisted of perturbations whose viability signatures closely resembled the global viability signature, and was similarly enriched for genes related to the regulation of cell cycle and apoptosis (Figure 5B). This result suggests that this cluster of perturbations is driving the global viability signature described above.

Interestingly, our analysis also revealed a smaller cluster of perturbations with a distinct transcriptional signature from the global viability signature. In fact, several of the perturbations in this group — such as the compound BAX-channel-blocker, and knockdown of genes such as FGFR3 and ATG5 — showed expression-viability relationships that were anticorrelated with the global viability signature and the signatures of compounds in the predominant cluster. Indeed, upon closer inspection, these perturbations seemed to be selectively inducing increased cell growth in certain cell lines, and these cell lines with increased proliferation showed a stronger transcriptional response compared to the other cell lines (Figure 5E). This result was further supported by enrichment for proliferation-related genes in these perturbations’ viability-related signatures. Several of the compounds and genetic perturbations in this cluster are thought to induce cell proliferation based on previous studies. For example, the small molecule BAX-channel-blocker inhibits cytochrome C release, preventing apoptosis [9]. In addition, decreased expression of FGFR3 and ATG5 have been shown to be associated with increased proliferation [13,19]. The fact that this set of perturbations elicited a transcriptional response that was partially overlapping with the larger set of perturbations that induced a cell cycle and apoptosis response suggests that similar cellular pathways might be implicated after treatment with both cell death- and proliferation-inducing perturbations, although they could be modulated differently.

These two distinct clusters, along with the wide variety of other perturbation-specific signatures, highlight the heterogeneity that exists in expression programs induced by selective treatments, and the advantages of analyzing the relationship between expression and viability on a perturbation-specific level.

## Discussion

We present an analysis of post-perturbational gene expression patterns that are associated with long-term viability in cancer cell lines. Applying global analyses across perturbations and cell lines, we found a robust relationship between transcriptional response and cell viability that was consistent across perturbation types (chemical and genetic), time-points, and dependency datasets. When applying more detailed analyses on a perturbation-specific level, however, we found that individual chemical and genetic reagents elicit a heterogeneous set of relationships between transcriptional change and cell viability.

Analysis of perturbation-specific viability signatures revealed several distinct clusters, most notably two broad sets of perturbations with anti-correlated viability signatures. These perturbation classes appear to have opposing effects on cell viability, yet produce similar patterns of transcriptional response in sensitive cells. Further examination of the cluster of perturbations that appeared to induce selective increases of cell proliferation suggested that they could be modulating cellular pathways in a way that promotes cell growth. Indeed, previous studies have found evidence that several of the compounds stimulate growth [9,12,13].

Surprisingly, this cluster of apparently proliferation-inducing perturbations produced transcriptional responses that were largely overlapping with that of perturbations in the predominant cluster. Given that the ‘global viability signature’ was characterized by activation of apoptosis and cell cycle arrest responses, this suggests that shared pathways of cellular stress may be involved in mediating responses to perturbations that result in increased growth as well as loss of cell viability, and that some genes in these pathways are modulated upward or downward regardless of the growth phenotype caused by a compound.

Our findings also suggest that analyzing the underlying components of the transcriptional drug response (viability-related and -independent) offers a more detailed view of the perturbation effect than traditional approaches. One approach commonly employed in the analysis of L1000 signatures is to compute their connectivity within each cell context individually, and then to summarize those connectivities across cell lines to derive and additional aggregate connectivity metric. This can be useful for identifying relationships that persist across multiple contexts or when it is unclear a priori which context to consider. However, it is not guaranteed to capture cell-context-specific effects, and only a portion of these effects are likely to be associated with the resulting cell death or cell cycle arrest processes. By integrating measured viability data, our modeling approach explicitly decomposes the transcriptional response to a perturbation into a viability-related component (which could suggest a drug’s killing mechanism), and a viability-independent component (which could better resolve the drug’s MOA, as was recently shown [24].

While preparing this manuscript, a similar analysis was reported investigating the patterns of association between expression and viability using several of the same datasets [24]. The authors’ results are largely concordant with our findings. In particular, both studies find the transcription-viability relationship to be robust across perturbation types and time points, and both demonstrate the ability to predict long-term sensitivity from post-perturbation expression profiles. Our approach extends their analysis by incorporating the recently published PRISM Repurposing drug sensitivity data [3], which allowed us to more thoroughly examine the expression-viability associations that are specific to individual perturbations. As demonstrated in the Results section, these individual signatures are crucial not only for understanding the different mechanisms underlying viability effects, but also for the development of pharmacodynamic markers of response that could be used in clinical applications.

Although our analysis focused on compounds that were screened in a sufficient number of cell lines, a limitation of the broader L1000 dataset is the small number of cell lines in which most compounds were profiled. With a larger panel of cell lines, analyses would be more powered to identify more subtle transcriptional responses that are specific to sensitive cell lines. Future studies will benefit from screening a large, consistent panel of samples across a diverse drug library. Indeed, single-cell sequencing technology has already proven useful for efficiently profiling many cell lines [16,26,10], and one study in particular found similar associations with drug sensitivity to those reported here [16].

Lastly, our findings support the feasibility of using rapid assays measuring the transcriptional response to infer the perturbation’s long-term effects on viability. This paradigm could be extended to the clinical realm by measuring expression responses of primary tumor cells to different drug treatments ex vivo in order to identify the most effective therapy for a patient. Indeed, because our results suggest that measuring transcriptional response even as early as 6 hours after treatment could yield useful predictions, this approach could overcome many of the current challenges of measuring drug response directly through ex vivo tumor cultures, such as obtaining a sufficient cell input and long-term cell culturing. The potential for this type of clinical application, along with our results demonstrating strong expression-viability relationships across many classes of drugs, suggest that the integration of these two data types could be useful for a wide array of uses.

## Methods

### Data

#### L1000 expression data

L1000 expression data (processed at “Level 5”) for compounds, shRNA, and **CRISPR** were downloaded from clue.io in the form of GCTx files [7]. The L1000 assay directly measures the expression of 978 landmark genes, and allows for computational estimation of about 12,000 more genes [23]. We use all genes (landmark and non-landmark) for most analyses, but subset to landmark genes for predictive modeling.

#### PRISM viability data

Drug response data were collected by the PRISM Repurposing project at the Broad Institute [3], and were downloaded from the Cancer Dependency Map portal https://depmap.org/portal/download/. The 19Q3 PRISM Repurposing Primary Screen data were used. PRISM is a high-throughput drug sensitivity assay that uses pools of barcoded cell lines [28]. Cell viability measurements were taken 5 days after initial drug treatment. GDSC viability data

The drug response data from the Wellcome Sanger Institute were downloaded from the Genomics of Drug Sensitivity in Cancer portal: https://www.cancerrxgene.org/downloads. The processed dose-response curve AUC values were used for defining a viability signature. Cell viability measurements were taken 3 days after initial drug treatment. CTD2 viability data

The drug response data from the Cancer Target Discovery and Development group were downloaded from the Cancer Therapeutics Response Portal: https://ocg.cancer.gov/programs/ctd2/data-portal. Cell viability measurements were taken 3 days after initial drug treatment. The results published here are fully or partially based upon data generated by the Cancer Target Discovery and Development (CTD2) Network (https://ocg.cancer.gov/programs/ctd2/data-portal) established by the National Cancer Institute’s Office of Cancer Genomics.

#### Normalizing viability datasets

In order to account for discrepancies between the PRISM, GDSC, and CTD2 datasets, we applied quantile normalization (learning the normalizing transformation from the overlapping data). Each dataset measures sensitivity as the area under the curve (AUC) of the dose-response curve. For interpretability, we transformed the datasets so that a value of 0 indicates no change in sensitivity, i.e., we used 1 - AUC as the sensitivity for all analyses.

#### RNAi viability data

The RNAi viability data were downloaded from the Cancer Dependency Map portal (https://depmap.org/portal/download/). The “gene means proc” scores from the 19Q2 data were used, which were estimated using DEMETER2 [15,6]. The sign of the scores were flipped to match the directionality of the drug sensitivity data.

#### CRISPR viability data

The CRISPR viability data were downloaded from the Cancer Dependency Map portal (https://depmap.org/portal/download/). The “gene effect corrected” scores from the 19Q1 Avana data were used [5,4].

### Statistical analysis

#### TAS

The transcriptional activity score (TAS), developed by the Connectivity Map, serves as a measure of the overall magnitude and consistency of a level-5 L1000 expression profile [23]. It is defined as:

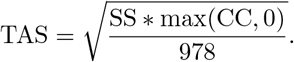

where SS is the signature strength and CC is the correlation between replicates. See [23] for more details.

#### Global viability signature

To compute the global viability signature, we pooled the data from all compounds, cell lines, and doses, and fit a linear model, regressing each gene’s expression on the matched viability values. Specifically, for each gene *i*, we fit the model

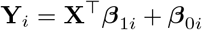

across all *n* profiles, where **X** ∊ *R*^*n*^ is the measured sensitivity values and **Y**_*i*_ ∊ *R*^*n*^ is a vector containing the level 5 L1000 differential expression values of gene *i* across experimental conditions. In the global analysis, the coefficients *β*_1*i*_ were extracted and used as the global viability-related signature coefficient for gene *i*. We did not make use of the *β*_0*i*_ coefficients for the global analysis and returned to them in the per-compound analysis. We fit these models using the R software package limma, which computes moderated t-statistics by shrinking gene-wise variance estimates to a common distribution computed across genes.

While the L1000 assay measures the expression of only 978 genes directly, the expression levels of about 12,000 genes are computationally inferred from these direct measurements. We included all genes (landmark and inferred) when computing the global viability signature.

#### Compound-specific viability signatures

To compute compound-specific viability signatures, we used a similar modeling procedure as above: for each compound *j* and each gene *i*, we fit the model

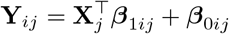

where **Y**_*ij*_ is a vector gene *i*’s differential expression following treatment with compound *j*, and **X**_*j*_ is a vector of measured sensitivity across cell lines following treatment. For each compound, we extract the ***β***_1*ij*_ coefficients as the “viability-related” signature and the ***β***_0*ij*_ coefficients as the “viability-independent” signatures for compound *j*. Because a sensitivity value of 0 corresponds to no viability effect, the intercepts ***β***_0*ij*_ correspond to the estimate of expression change in cell line that exhibits no sensitivity effects.

Our model fits a univariate linear model for each gene separately, so it is agnostic to any interaction or collinearity that may exist between the expression of different genes. Compared to a model that considers all genes as covariates simultaneously, we find that this approach allows for a simpler interpretation of the gene-wise coefficients. However, we use this alternative approach when predicting sensitivity from expression changes, as described below.

#### Predictive modeling

To predict long-term sensitivity using short-term expression across all compounds, we fit a linear model with the 978 landmark gene expression values as covariates and the measured sensitivity as the response variable. We used elastic net linear regression, which applies a combination of *L*1 and *L*2 regularization. To measure the accuracy of the model’s predictions, we used a 10-fold cross-validation procedure (where the splits were based on compounds), and measured the Pearson correlation between the model’s predicted sensitivity values and the true sensitivity for each test split.

To assess the predictivity of compound-specific differential sensitivity across cell lines, we again used elastic net linear regression, but assessed the model with a leave-one-compound-out procedure, training on *n*–1 compounds (where *n* is the total number of compounds), and testing on the one held-out compound. We again used the Pearson correlation between the predicted and true sensitivity as a measure of the accuracy of the model.

Finally, to assess the between-compound variation in expression to the previous models’ performance, we fit a linear model using a reduced version of each compound’s transcriptional data. We averaged each compound’s original profiles, yielding a single 978-length expression vector for each compound. Hence, the model did not have access to any information about each compound’s variation in expression patterns across cell lines, and was forced to rely on the expression variation across compounds in order to predict sensitivity.

#### Gene set enrichment analysis

Normalized gene set enrichment scores, and asociated p-values, were computed by a gene permutation-based procedure using the R package fgsea with the MSigDB Hallmark gene set collection.

#### Viability signature clustering analysis

Perturbation-specific viability signatures were clustered and displayed using the R pheatmap package. Clusters of perturbation signatures were further identified and refined using hierarchical clustering via the R function hclust.

#### Agreement between sensitivity datasets

In order to validate the robustness of these viability signatures, we computed the signatures separately for each of the three large-scale drug sensitivity datasets (PRISM, GDSC, and CTRP). The expression-viability relationship remained consistent across the three screens, reinforcing the robustness of this association (Figure S1). Given this broad agreement, we use the normalized average of the three datasets for the remaining analyses (Methods). Interpreting viability signature components

Examining the magnitude and direction of the viability-related and -independent components for a compound can help characterize its viability signature. For example, a compound with a zero-magnitude viability-independent component but a strong nonzero-magnitude viability-related component indicates that the expression induced by the compound is entirely associated with loss of viability. Alternatively, suppose a compound has nonzero-magnitude viability-independent component -related components, and these two components have different directions. This would suggest that the compound induces one transcriptional program in all cells (regardless of sensitivity), and another program that is specifically expressed in sensitive cells.

## Code availability

All code for analyses and figures in this manuscript is available at https://github.com/andrewcharlesjones/l1000_viability_signature_manuscript. Custom code for general L1000 data formatting and analysis is available at https://github.com/andrewcharlesjones/l1000_analysis.

## Acknowledgments

The authors would like to thank Todd Golub, Aravind Subramanian, Ted Natoli, Rajiv Narayan, and Mustafa Kocak for helpful comments and suggestions.

## Supplementary Material

**Fig. S1:**
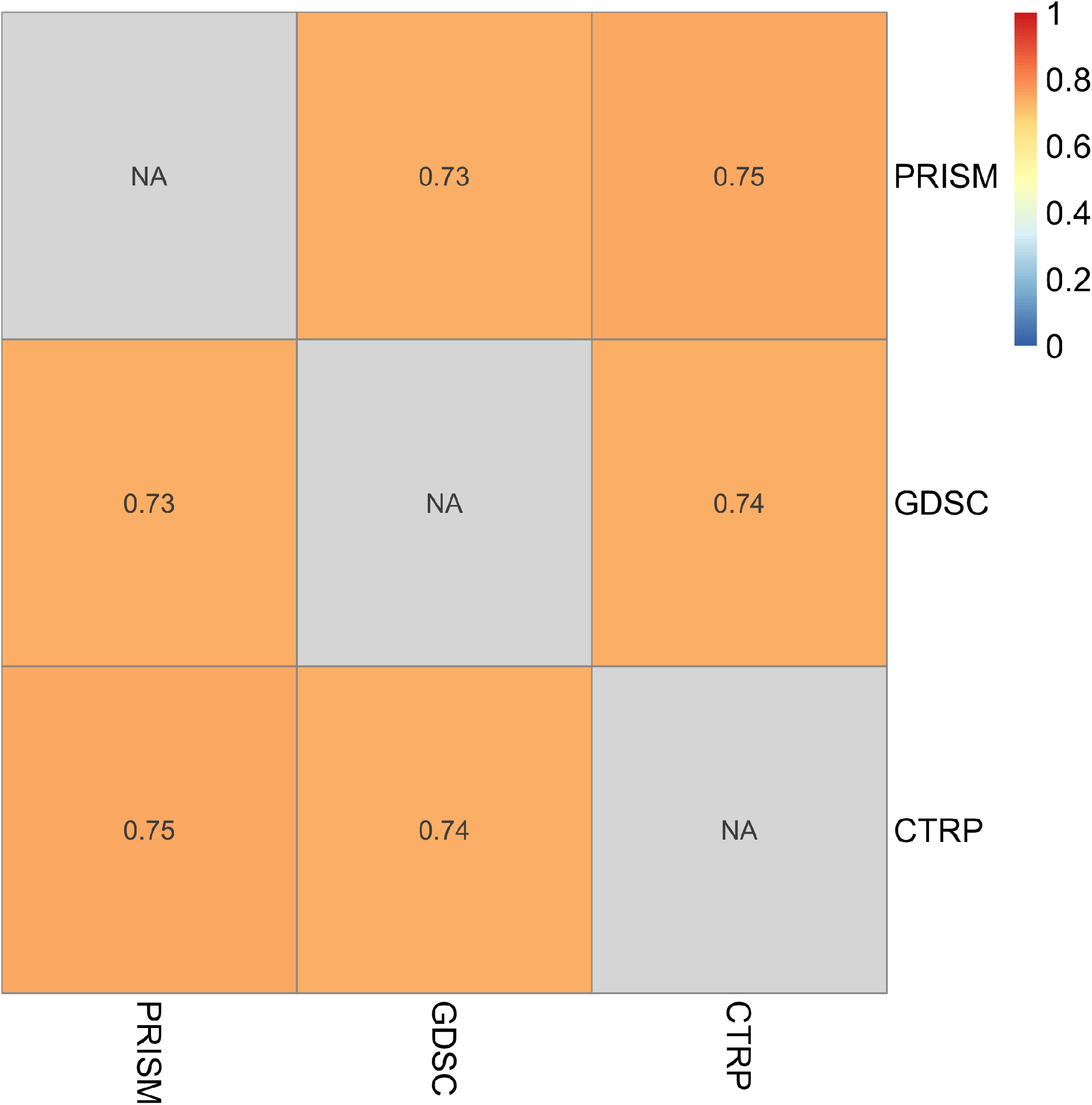
Agreement between global viability signature across three independent datasets. Strong agreement was found between the transcription-viability relationship, as estimated on three independent drug sensitivity datasets: PRISM, GDSC, and CTRP (see Methods). Each box shows the pairwise Pearson correlation between each viability signature.

**Fig. S2:**
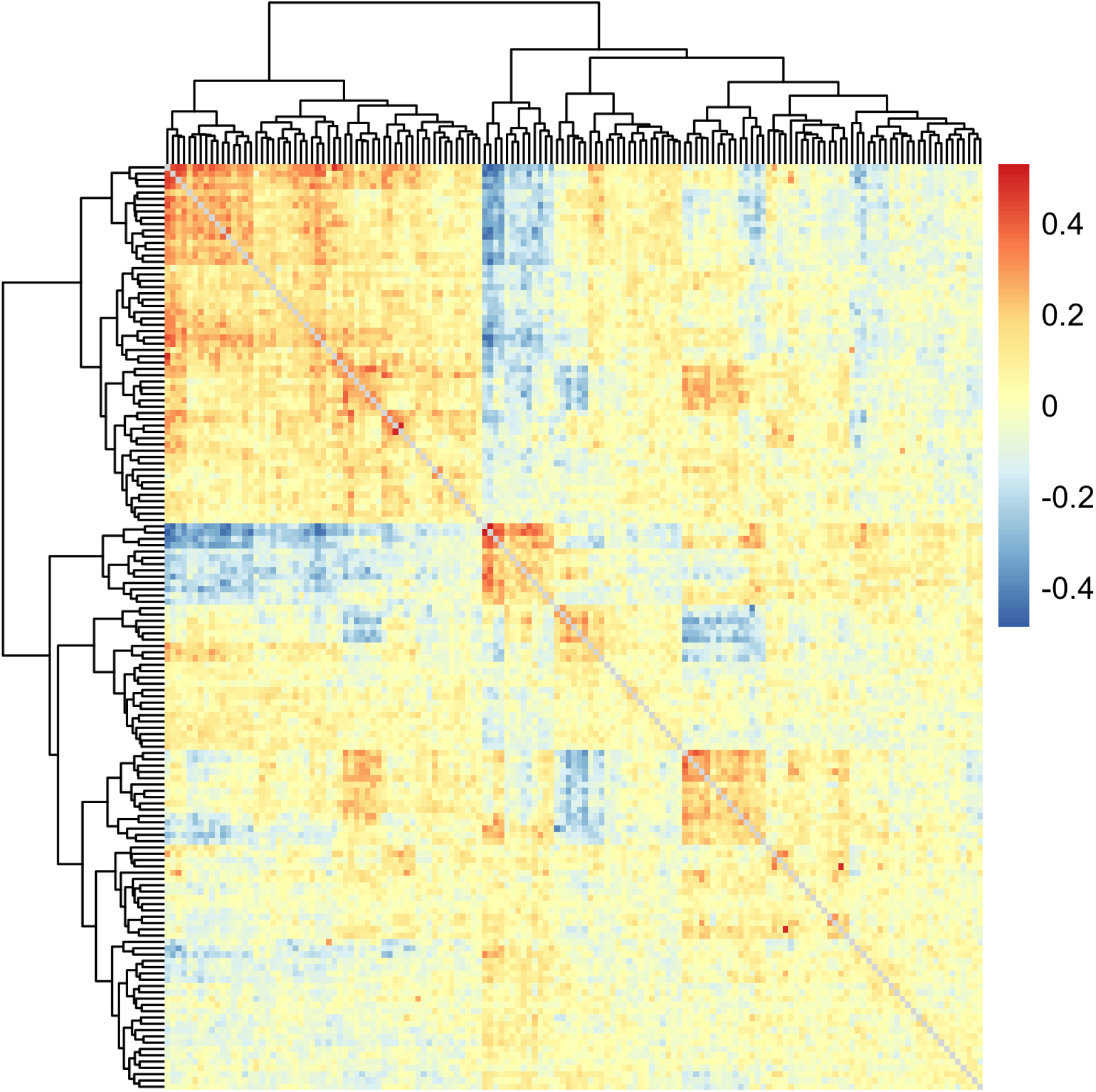
Perturbation-specific viability signatures showed substantial heterogeneity, as well as tight clustering among some subsets of drugs. Perturbation-specific signatures represent the transcription-viability signature uniquely for each compound. Shown above are the pairwise Pearson cor-relations between all drug-specific signatures, showing that clear differences exist among the signatures, but some drugs elicit similar transcriptional patterns in relation to sensitivity. Drug labels on the axes are omitted for clarity, but see Figure S3 for a smaller example.

**Fig. S3:**
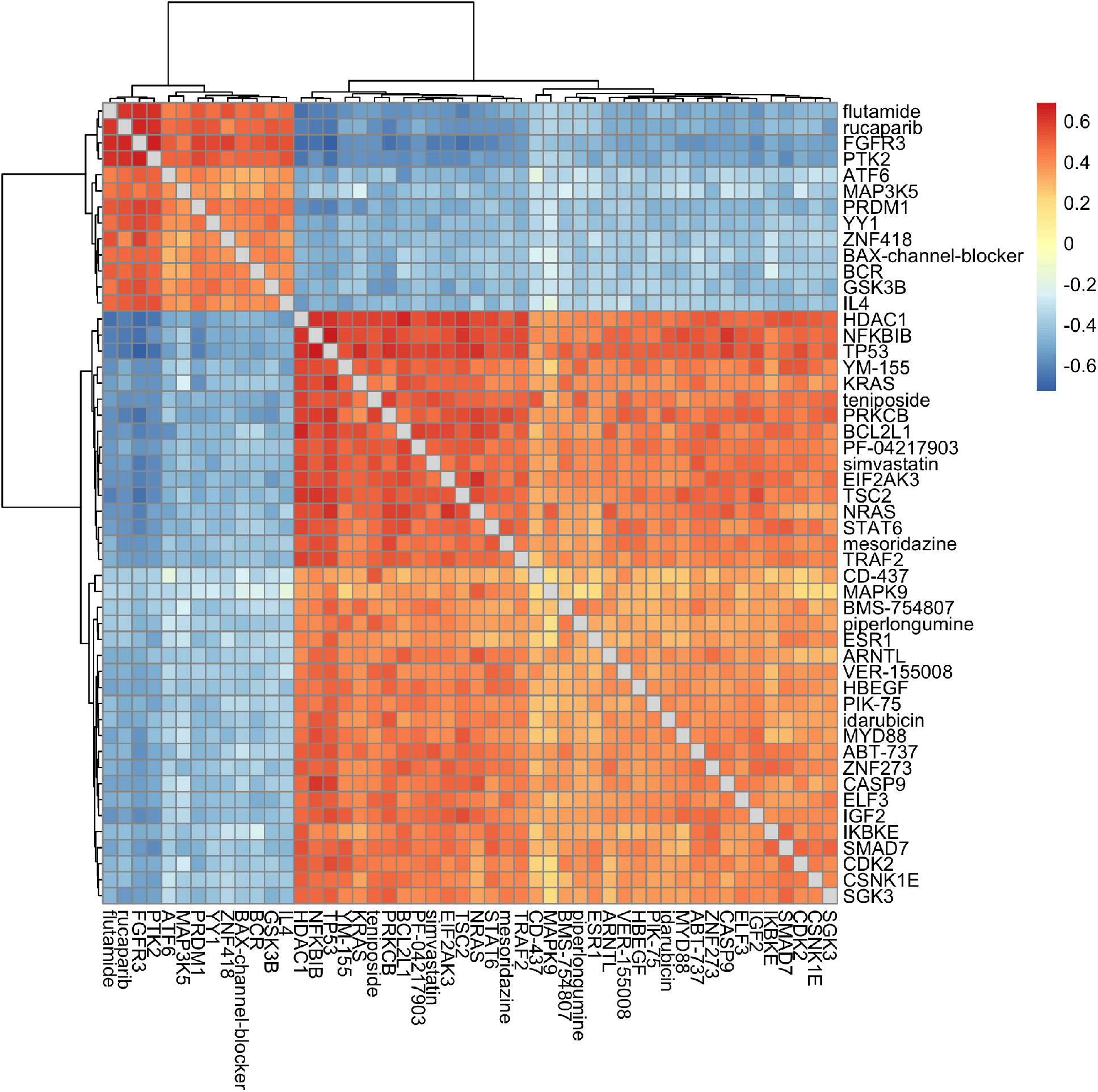
Two clusters of drugs with anticorrelated viability signatures. Shown here are the Pearson correlations between perturbation-specific viability signatures for a subset of perturbations (compounds and shRNA, in this case) showing the contrast between the two identified clusters.

## References

1. Basu, A., Bodycombe, N.E., Cheah, J.H., Price, E.V., Liu, K., Schaefer, G.I., Ebright, R.Y., Stewart, M.L., Ito, D., Wang, S., Bracha, A.L., Liefeld, T., Wawer, M., Gilbert, J.C., Wilson, A.J., Stransky, N., Kryukov, G.V., Dancik, V., Barretina, J., Garraway, L.A., Hon, C.S.Y., Munoz, B., Bittker, J.A., Stockwell, B.R., Khabele, D., Stern, A.M., Clemons, P.A., Shamji, A.F., Schreiber, S.L.: An interactive resource to identify cancer genetic and lineage dependencies targeted by small molecules. Cell 154(5), 1151–1161 (aug 2013). https://doi.org/10.1016/j.cell.2013.08.003, http://dx.doi.org/10.1016/j.cell.2013.08.003

2. Corsello, S.M., Bittker, J.A., Liu, Z., Gould, J., McCarren, P., Hirschman, J.E., Johnston, S.E., Vrcic, A., Wong, B., Khan, M., Asiedu, J., Narayan, R., Mader, C.C., Subramanian, A., Golub, T.R.: The drug repurposing hub: a next-generation drug library and information resource. Nature Medicine 23(4), 405–408 (apr 2017). https://doi.org/10.1038/nm.4306, http://dx.doi.org/10.1038/nm.4306

3. Corsello, S.M., Nagari, R.T., Spangler, R.D., Rossen, J., Kocak, M., Bryan, J.G., Humeidi, R., Peck, D., Wu, X., Tang, A.A., Wang, V.M., Bender, S.A., Lemire, E., Narayan, R., Montgomery, P., Ben-David, U., Garvie, C.W., Chen, Y., Rees, M.G., Lyons, N.J., McFarland, J.M., Wong, B.T., Wang, L., Dumont, N., O, P.J., Stefan, E., Doench, J.G., Harrington, C.N., Greulich, H., Meyerson, M., Vazquez, F., Subramanian, A., Roth, J.A., Bittker, J.A., Boehm, J.S., Mader, C.C., Tsherniak, A., Golub, T.R.: Discovering the anticancer potential of non-oncology drugs by systematic viability profiling. Nature Cancer (jan 2020). https://doi.org/10.1038/s43018-019-0018-6, http://www.nature.com/articles/s43018-019-0018-6

4. Dempster, J.M., Rossen, J., Kazachkova, M., Pan, J., Kugener, G., Root, D.E., Tsherniak, A.: Extracting biological insights from the project achilles genome-scale CRISPR screens in cancer cell lines. BioRxiv (jul 2019). https://doi.org/10.1101/720243, http://biorxiv.org/lookup/doi/10.1101/720243

5. DepMap, B.: DEMETER 2 combined RNAi. Figshare (2019). https://doi.org/10.6084/m9.figshare.9170975.v1, http://figshare.com/articles/{DEMETER_2_Combined_RNAi}/9170975/1

6. DepMap, B.: DepMap achilles 19Q1 public. Figshare (2019). https://doi.org/10.6084/m9.figshare.7655150, http://figshare.com/articles/{DepMap_Achilles_19Q1_Public}/7655150

7. Enache, O.M., Lahr, D.L., Natoli, T.E., Litichevskiy, L., Wadden, D., Flynn, C., Gould, J., Asiedu, J.K., Narayan, R., Subramanian, A.: The GCTx format and cmap{Py, R, M} packages: resources for the optimized storage and integrated traversal of dense matrices of data and annotations. BioRxiv (nov 2017). https://doi.org/10.1101/227041, http://biorxiv.org/lookup/doi/10.1101/227041

8. Garnett, M.J., Edelman, E.J., Heidorn, S.J., Greenman, C.D., Dastur, A., Lau, K.W., Greninger, P., Thompson, I.R., Luo, X., Soares, J., Liu, Q., Iorio, F., Surdez, D., Chen, L., Milano, R.J., Bignell, G.R., Tam, A.T., Davies, H., Stevenson, J.A., Barthorpe, S., Lutz, S.R., Kogera, F., Lawrence, K., McLaren-Douglas, A., Mitropoulos, X., Mironenko, T., Thi, H., Richardson, L., Zhou, W., Jewitt, F., Zhang, T., O’Brien, P., Boisvert, J.L., Price, S., Hur, W., Yang, W., Deng, X., Butler, A., Choi, H.G., Chang, J.W., Baselga, J., Stamenkovic, I., Engelman, J.A., Sharma, S.V., Delattre, O., Saez-Rodriguez, J., Gray, N.S., Settleman, J., Futreal, P.A., Haber, D.A., Stratton, M.R., Ramaswamy, S., McDermott, U., Benes, C.H.: Systematic identification of genomic markers of drug sensitivity in cancer cells. Nature 483(7391), 570–575 (mar 2012). https://doi.org/10.1038/nature11005, http://dx.doi.org/10.1038/nature11005

9. Hetz, C., Vitte, P.A., Bombrun, A., Rostovtseva, T.K., Montessuit, S., Hiver, A., Schwarz, M.K., Church, D.J., Korsmeyer, S.J., Martinou, J.C., Antonsson, B.: Bax channel inhibitors prevent mitochondrion-mediated apoptosis and protect neurons in a model of global brain ischemia. The Journal of Biological Chemistry 280(52), 42960–42970 (dec 2005). https://doi.org/10.1074/jbc.M505843200, http://dx.doi.org/10.1074/jbc.M505843200

10. Huang, Y., McCarthy, D.J., Stegle, O.: Vireo: Bayesian demultiplexing of pooled single-cell RNA-seq data without genotype reference. BioRxiv (apr 2019). https://doi.org/10.1101/598748, http://biorxiv.org/lookup/doi/10.1101/598748

11. Iorio, F., Knijnenburg, T.A., Vis, D.J., Bignell, G.R., Menden, M.P., Schubert, M., Aben, N., Gonçalves, E., Barthorpe, S., Lightfoot, H., Cokelaer, T., Greninger, P., van Dyk, E., Chang, H., de Silva, H., Heyn, H., Deng, X., Egan, R.K., Liu, Q., Mironenko, T., Mitropoulos, X., Richardson, L., Wang, J., Zhang, T., Moran, S., Sayols, S., Soleimani, M., Tamborero, D., Lopez-Bigas, N., Ross-Macdonald, P., Esteller, M., Gray, N.S., Haber, D.A., Stratton, M.R., Benes, C.H., Wessels, L.F.A., Saez-Rodriguez, J., McDermott, U., Garnett, M.J.: A landscape of pharmacogenomic interactions in cancer. Cell 166(3), 740–754 (jul 2016). https://doi.org/10.1016/j.cell.2016.06.017, http://linkinghub.elsevier.com/retrieve/pii/S0092867416307462

12. Juürgensmeier, J.M., Xie, Z., Deveraux, Q., Ellerby, L., Bredesen, D., Reed, J.C.: Bax directly induces release of cytochrome c from isolated mitochondria. Proceedings of the National Academy of Sciences of the United States of America 95(9), 4997–5002 (apr 1998). https://doi.org/10.1073/pnas.95.9.4997, http://dx.doi.org/10.1073/pnas.95.9.4997

13. Lafitte, M., Moranvillier, I., Garcia, S., Peuchant, E., Iovanna, J., Rousseau, B., Dubus, P., Guyonnet-Dupérat, V., Belleannée, G., Ramos, J., Bedel, A., de Verneuil, H., Moreau-Gaudry, F., Dabernat, S.: FGFR3 has tumor suppressor properties in cells with epithelial phenotype. Molecular Cancer 12, 83 (jul 2013). https://doi.org/10.1186/1476-4598-12-83, http://dx.doi.org/10.1186/1476-4598-12-83

14. Lamb, J., Crawford, E.D., Peck, D., Modell, J.W., Blat, I.C., Wrobel, M.J., Lerner, J., Brunet, J.P., Subramanian, A., Ross, K.N., Reich, M., Hieronymus, H., Wei, G., Armstrong, S.A., Haggarty, S.J., Clemons, P.A., Wei, R., Carr, S.A., Lander, E.S., Golub, T.R.: The connectivity map: using gene-expression signatures to connect small molecules, genes, and disease. Science 313(5795), 1929–1935 (sep 2006). https://doi.org/10.1126/science.1132939, http://dx.doi.org/10.1126/science.1132939

15. McFarland, J.M., Ho, Z.V., Kugener, G., Dempster, J.M., Montgomery, P.G., Bryan, J.G., Krill-Burger, J.M., Green, T.M., Vazquez, F., Boehm, J.S., Golub, T.R., Hahn, W.C., Root, D.E., Tsherniak, A.: Improved estimation of cancer dependencies from large-scale RNAi screens using model-based normalization and data integration. Nature Communications 9(1), 4610 (nov 2018). https://doi.org/10.1038/s41467-018-06916-5, http://www.nature.com/articles/s41467-018-06916-5

16. McFarland, J.M., Paolella, B.R., Warren, A., Geiger-Schuller, K., Shibue, T., Rothberg, M., Kuksenko, O., Jones, A., Chambers, E., Dionne, D., Bender, S., Wolpin, B.M., Ghandi, M., Tirosh, I., Rozenblatt-Rosen, O., Roth, J.A., Golub, T.R., Regev, A., Aguirre, A.J., Vazquez, F., Tsherniak, A.: Multiplexed single-cell profiling of post-perturbation transcriptional responses to define cancer vulnerabilities and therapeutic mechanism of action. BioRxiv (dec 2019). https://doi.org/10.1101/868752, http://biorxiv.org/lookup/doi/10.1101/868752

17. Meyers, R.M., Bryan, J.G., McFarland, J.M., Weir, B.A., Sizemore, A.E., Xu, H., Dharia, N.V., Montgomery, P.G., Cowley, G.S., Pantel, S., Goodale, A., Lee, Y., Ali, L.D., Jiang, G., Lubonja, R., Harrington, W.F., Strickland, M., Wu, T., Hawes, D.C., Zhivich, V.A., Wyatt, M.R., Kalani, Z., Chang, J.J., Okamoto, M., Stegmaier, K., Golub, T.R., Boehm, J.S., Vazquez, F., Root, D.E., Hahn, W.C., Tsherniak, A.: Computational correction of copy number effect improves specificity of CRISPR-cas9 essentiality screens in cancer cells. Nature Genetics 49(12), 1779–1784 (dec 2017). https://doi.org/10.1038/ng.3984, http://www.nature.com/doifinder/10.1038/ng.3984

18. Niepel, M., Hafner, M., Duan, Q., Wang, Z., Paull, E.O., Chung, M., Lu, X., Stuart, J.M., Golub, T.R., Subramanian, A., Ma’ayan, A., Sorger, P.K.: Common and cell-type specific responses to anti-cancer drugs revealed by high throughput transcript profiling. Nature Communications 8(1), 1186 (oct 2017). https://doi.org/10.1038/s41467-017-01383-w, http://dx.doi.org/10.1038/s41467-017-01383-w

19. Qin, Y., Sun, W., Zhang, H., Zhang, P., Wang, Z., Dong, W., He, L., Zhang, T., Shao, L., Zhang, W., Wu, C.: LncRNA GAS8-AS1 inhibits cell proliferation through ATG5-mediated autophagy in papillary thyroid cancer. Endocrine 59(3), 555–564 (jan 2018). https://doi.org/10.1007/s12020-017-1520-1, http://dx.doi.org/10.1007/s12020-017-1520-1

20. Rampášek, L., Hidru, D., Smirnov, P., Haibe-Kains, B., Goldenberg, A.: Dr.VAE: improving drug response prediction via modeling of drug perturbation effects. Bioinformatics 35(19), 3743–3751 (oct 2019). https://doi.org/10.1093/bioinformatics/btz158, http://dx.doi.org/10.1093/bioinformatics/btz158

21. Rees, M.G., Seashore-Ludlow, B., Cheah, J.H., Adams, D.J., Price, E.V., Gill, S., Javaid, S., Coletti, M.E., Jones, V.L., Bodycombe, N.E., Soule, C.K., Alexander, B., Li, A., Montgomery, P., Kotz, J.D., Hon, C.S.Y., Munoz, B., Liefeld, T., Dančík, V., Haber, D.A., Clish, C.B., Bittker, J.A., Palmer, M., Wagner, B.K., Clemons, P.A., Shamji, A.F., Schreiber, S.L.: Correlating chemical sensitivity and basal gene expression reveals mechanism of action. Nature Chemical Biology 12(2), 109–116 (feb 2016). https://doi.org/10.1038/nchembio.1986, http://dx.doi.org/10.1038/nchembio.1986

22. Seashore-Ludlow, B., Rees, M.G., Cheah, J.H., Cokol, M., Price, E.V., Coletti, M.E., Jones, V., Bodycombe, N.E., Soule, C.K., Gould, J., Alexander, B., Li, A., Montgomery, P., Wawer, M.J., Kuru, N., Kotz, J.D., Hon, C.S.Y., Munoz, B., Liefeld, T., Dančík, V., Bittker, J.A., Palmer, M., Bradner, J.E., Shamji, A.F., Clemons, P.A., Schreiber, S.L.: Harnessing connectivity in a large-scale small-molecule sensitivity dataset. Cancer discovery 5(11), 1210–1223 (nov 2015). https://doi.org/10.1158/2159-8290.CD-15-0235, http://dx.doi.org/10.1158/2159-8290.{CD}-15-0235

23. Subramanian, A., Narayan, R., Corsello, S.M., Peck, D.D., Natoli, T.E., Lu, X., Gould, J., Davis, J.F., Tubelli, A.A., Asiedu, J.K., Lahr, D.L., Hirschman, J.E., Liu, Z., Donahue, M., Julian, B., Khan, M., Wadden, D., Smith, I.C., Lam, D., Liberzon, A., Toder, C., Bagul, M., Orzechowski, M., Enache, O.M., Piccioni, F., Johnson, S.A., Lyons, N.J., Berger, A.H., Shamji, A.F., Brooks, A.N., Vrcic, A., Flynn, C., Rosains, J., Takeda, D.Y., Hu, R., Davison, D., Lamb, J., Ardlie, K., Hogstrom, L., Greenside, P., Gray, N.S., Clemons, P.A., Silver, S., Wu, X., Zhao, W.N., Read-Button, W., Wu, X., Haggarty, S.J., Ronco, L.V., Boehm, J.S., Schreiber, S.L., Doench, J.G., Bittker, J.A., Root, D.E., Wong, B., Golub, T.R.: A next generation connectivity map: L1000 platform and the first 1,000,000 profiles. Cell 171(6), 1437–1452.e17 (nov 2017). https://doi.org/10.1016/j.cell.2017.10.049, http://linkinghub.elsevier.com/retrieve/pii/S0092867417313090

24. Szalai, B., Subramanian, V., Holland, C.H., Alföldi, R., Puskás, L.G., Saez-Rodriguez, J.: Signatures of cell death and proliferation in perturbation transcriptomics data-from confounding factor to effective prediction. Nucleic Acids Research 47(19), 10010–10026 (nov 2019). https://doi.org/10.1093/nar/gkz805, http://dx.doi.org/10.1093/nar/gkz805

25. Tsherniak, A., Vazquez, F., Montgomery, P.G., Weir, B.A., Kryukov, G., Cowley, G.S., Gill, S., Harrington, W.F., Pantel, S., Krill-Burger, J.M., Meyers, R.M., Ali, L., Goodale, A., Lee, Y., Jiang, G., Hsiao, J., Gerath, W.F.J., Howell, S., Merkel, E., Ghandi, M., Garraway, L.A., Root, D.E., Golub, T.R., Boehm, J.S., Hahn, W.C.: Defining a cancer dependency map. Cell 170(3), 564–576.e16 (jul 2017). https://doi.org/10.1016/j.cell.2017.06.010, http://linkinghub.elsevier.com/retrieve/pii/S0092867417306517

26. Xu, J., Falconer, C., Coin, L.: Genotype-free demultiplexing of pooled single-cell RNA-seq. BioRxiv (mar 2019). https://doi.org/10.1101/570614, http://biorxiv.org/lookup/doi/10.1101/570614

27. Yang, W., Soares, J., Greninger, P., Edelman, E.J., Lightfoot, H., Forbes, S., Bindal, N., Beare, D., Smith, J.A., Thompson, I.R., Ramaswamy, S., Futreal, P.A., Haber, D.A., Stratton, M.R., Benes, C., McDermott, U., Garnett, M.J.: Genomics of drug sensitivity in cancer (GDSC): a resource for therapeutic biomarker discovery in cancer cells. Nucleic Acids Research 41(Database issue), D955–61 (jan 2013). https://doi.org/10.1093/nar/gks1111, http://dx.doi.org/10.1093/nar/gks1111

28. Yu, C., Mannan, A.M., Yvone, G.M., Ross, K.N., Zhang, Y.L., Marton, M.A., Taylor, B.R., Crenshaw, A., Gould, J.Z., Tamayo, P., Weir, B.A., Tsherniak, A., Wong, B., Garraway, L.A., Shamji, A.F., Palmer, M.A., Foley, M.A., Winckler, W., Schreiber, S.L., Kung, A.L., Golub, T.R.: High-throughput identification of genotype-specific cancer vulnerabilities in mixtures of barcoded tumor cell lines. Nature Biotechnology 34(4), 419–423 (apr 2016). https://doi.org/10.1038/nbt.3460, http://www.nature.com/doifinder/10.1038/nbt.3460

